# Use of a truffle dog provides insight into the ecology and abundant occurrence of *Genea* (Pyronemataceae) in western Oregon, U.S.A

**DOI:** 10.1101/2024.07.29.604843

**Authors:** Hilary Rose Dawson, Heather A. Dawson

**Author notes:** indicates authors contributed equally.

## Abstract

Hypogeous fungi (“truffles”) are challenging to study because they produce underground sporocarps that may not be found during traditional fungal surveys. Commercially valuable truffles are located using scent-detection dogs trained on truffles. However, the dogs are not necessarily limited to commercial truffle species when trained on other truffle taxa of interest. For example, *Genea* (Pyronemataceae, Ascomycota) is a genus of common but difficult to find truffles that develops small and often soil-colored hypogeous ascomata (truffles).. We used a truffle dog trained to locate *Genea* truffles in the southern Willamette Valley and associated mountains in western Oregon, U.S.A, recording when the sporocarps were present at a wide range of elevations (113 to 1879 m). We found *Genea* was present in half of our surveys, and noted that it rarely fruited in areas that had experienced wildfire. This study demonstrates the value of using truffle dogs in documenting truffle diversity, particularly those that are difficult to locate visually, and provides further evidence for the abundance of *Genea*.

## Introduction

The truffle genus *Genea* (Pyronemataceae, Ascomycota) is characterized by lobed and often inconspicuous hypogeous ascomata and was originally circumscribed in Italy in the 1800s (Vittadini, 1831). It is considered common in both Europe and North America (Alvarado et al., 2016), but because it fruits underground and is often difficult to differentiate from soil aggregates, we have limited knowledge of its ecology and distribution. One of the most comprehensive studies in the U.S. Pacific Northwest focused on oak grasslands (Smith et al., 2006) and described two new species. However, *Genea* forms ectomycorrhizal associations with a wide variety of plants (Smith et al., 2006), suggesting that it may be widely distributed throughout the Douglas-fir (*Pseudotsuga menziesii*) forests of the Cascades and Coast Range. We tested this possibility using a novel tool: a truffle dog with an unusual ability to find *Genea* truffles.

Scent-detection dogs are used to detect odors in a wide range of contexts such as military, medical, and increasingly in conservation (Beebe et al., 2016). In conservation, detection dogs are primarily used to detect animal species of interest, but are also used to detect plants, fungi, and even bacteria of interest (Grimm-Seyfarth et al., 2021). Commercially, scent-detection dogs have been used to locate culinary truffles (Tuber sp.) for centuries (Čejka et al., 2022)However, some dogs have a natural inclination towards generalizing on any hypogeous odorous sporocarps. If the trainer rewards all finds (rather than just culinarily valuable species), they can train “diversity dogs” who can locate a broad range of truffles that may be of interest to science. Given that detection dogs can be used to locate rare or otherwise difficult to find species (Bennett et al., 2020), detection dogs often outperform other methods (Grimm-Seyfarth et al., 2021), and some truffle species, like Genea, are visually cryptic but often highly aromatic. We set out to test if a such a “truffle diversity dog” who has demonstrated an aptitudefor Genea could increase our regional knowledge of this truffle. Traditionally, truffle surveys are often performed by raking, where a researcher uses a small garden hand rake to peel back the upper layer of the soil. This method has a visual bias towards what the researcher spots under the duff, and often misses rare and inconspicuous taxa. In contrast, truffle dogs have a scent bias driven by mature sporocarps of any size or color, can find truffles deeper than most people rake, and do not have preconceived ideas of where truffles ‘should’ be. In this study, we set out to document the ecology and abundance of Genea using a truffle dog, understanding that we were limited by the lack of taxonomic resolution in the genus in our region. Although Genea in the Pacific Northwest needs further systematic research, such analyses are outside the scope of this study. By documenting where a truffle genus is found throughout a range of sites and seasons, we can use truffle diversity dogs to broaden our understanding of hypogeous fungal ecology even without understanding the complexity of species present.

## Methods

All truffles were located by trained truffle dog Rye, who had found 23 genera of truffles in two years of truffling as of February 2023. Rye demonstrated particular persistence in locating *Genea* truffles as of the time of this study. He received a reward for each *Genea* truffle located, reinforcing his pursuit of truffles with this type of aroma.

We performed 87 general truffle surveys and forays in all seasons between October 2021 and November 2022. We visited 66 sites, 12 of which we visited more than once throughout the year (Figure 1; Table S1). Most surveys were conducted along existing public trails that allowed us easy access to varied landscapes. We performed these surveys at a range of elevations from 113 m to 1879 m (elevation data extracted from WorldClim (Fick & Hijmans, 2017)). On each survey, we primed Rye to find truffles of any species and rewarded his finds with praise and a brief game of fetch with a tennis ball (his favorite reward). He was given freedom to roam off-leash with voice recall during these surveys so that he could cover more ground and follow his nose.

**Figure 1:**
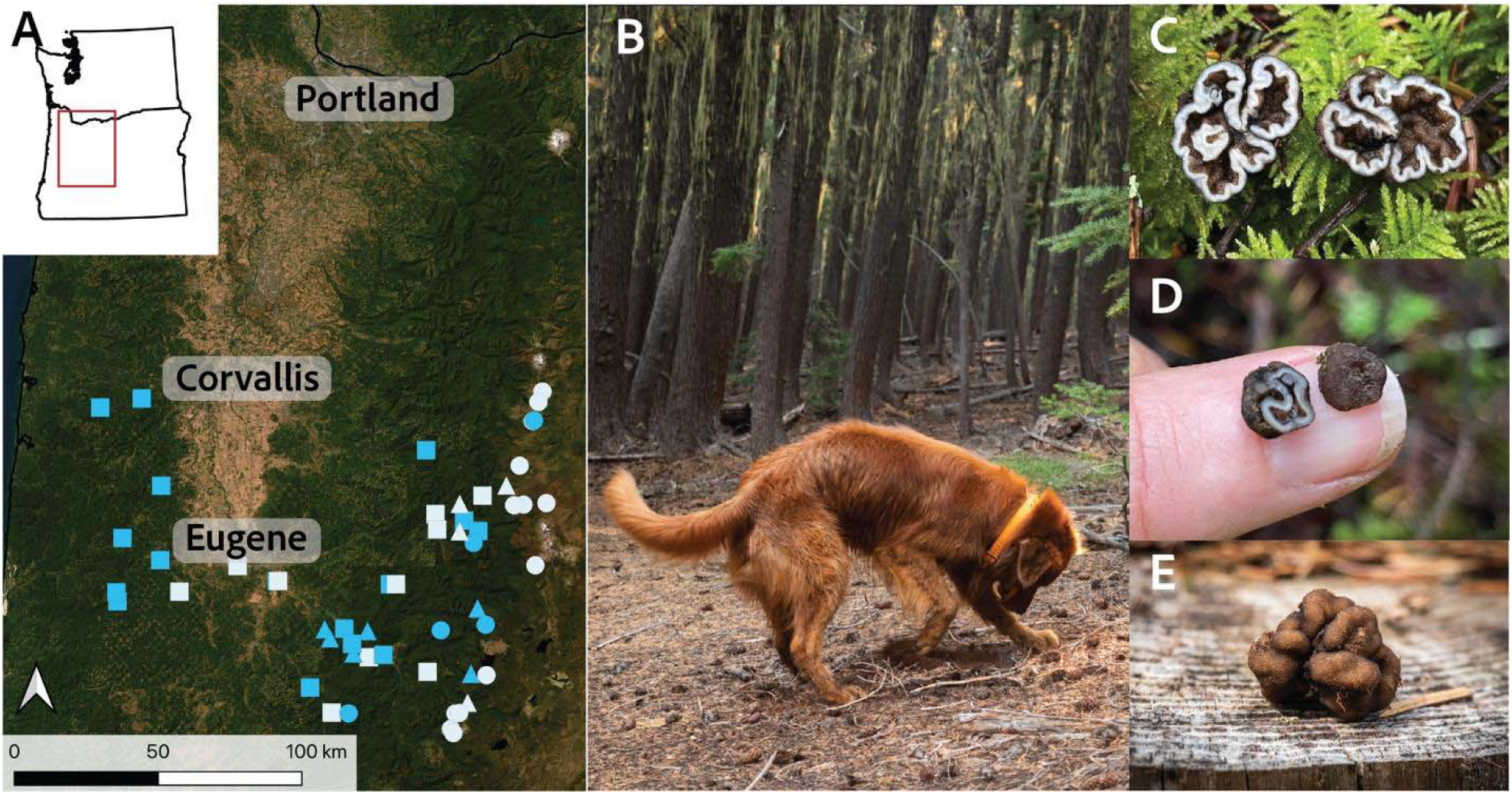
Sites and methods used to document *Genea* ecology. A) Map of all sites surveyed as part of this study. Blue points are sites where we found at least one *Genea* truffle. Symbols indicate elevation category (circle = high elevation, triangle = mid elevation, square = low elevation). B) Example of truffle dog Rye finding a truffle. C-E) Photographs of *Genea* truffles demonstrating lobing, dark peridium and light gleba, and sometimes very small size and resemblance to soil clusters.

We conducted our surveys primarily in conifer-dominated forests of varying age classes and densities, with the exception of the site at Mt. Pisgah which was oak savanna (Table S1). Our sites had a variety of fire histories, from no recent fire to recently burned (1-11 years). Most low-mid elevation conifer forest sites were primarily composed of Douglas-fir (*Pseudotsuga menziesii*) and western hemlock (*Tsuga heterophylla*), and high elevation sites often included mountain hemlock (*Tsuga mertensiana*), true fir (*Abies* spp.), and pine (*Pinus* spp.). All sites were located in Benton, Deschutes, Lane, and Linn Counties in southern Willamette Valley, Oregon, USA. These counties encompass the western Cascades, southern Coast Range, and Willamette Valley. We excluded observations from coastal Lane County because we performed only spatiotemporally limited surveys on the coast.

Genea was identified to genus level in the field based on macroscopic morphological features. Spores from a subset of our finds were checked to verify this identification. We only recorded truffle presence/absence at each site. Where multiple truffles were found, this counted as a single ‘presence’ for a site. If we surveyed multiple habitat types at a single location (for example, inside and outside a burn area), this was counted as multiple sites. Elevation was categorized as ‘low’ (0-750 m), ‘mid’ (750-1350 m), and ‘high’ (>1350 m) based on observed changes in plant communities and general ecology for the area.

## Results

We found *Genea* fruiting bodies in slightly over half (51.7%) of the site surveys. We reported occurrences of *Genea* nearly year-round, except the months of August and September (Figure 2) which are the driest months of the year due to western Oregon’s Mediterranean climate. Low and mid elevation sites had the most consistent *Genea* presence. The highest site we found *Genea* at was at 1716 m elevation. *Genea* truffles were not present in any burned conifer forests, but were found during two oak savanna surveys that had burned three years prior.

**Figure 2:**
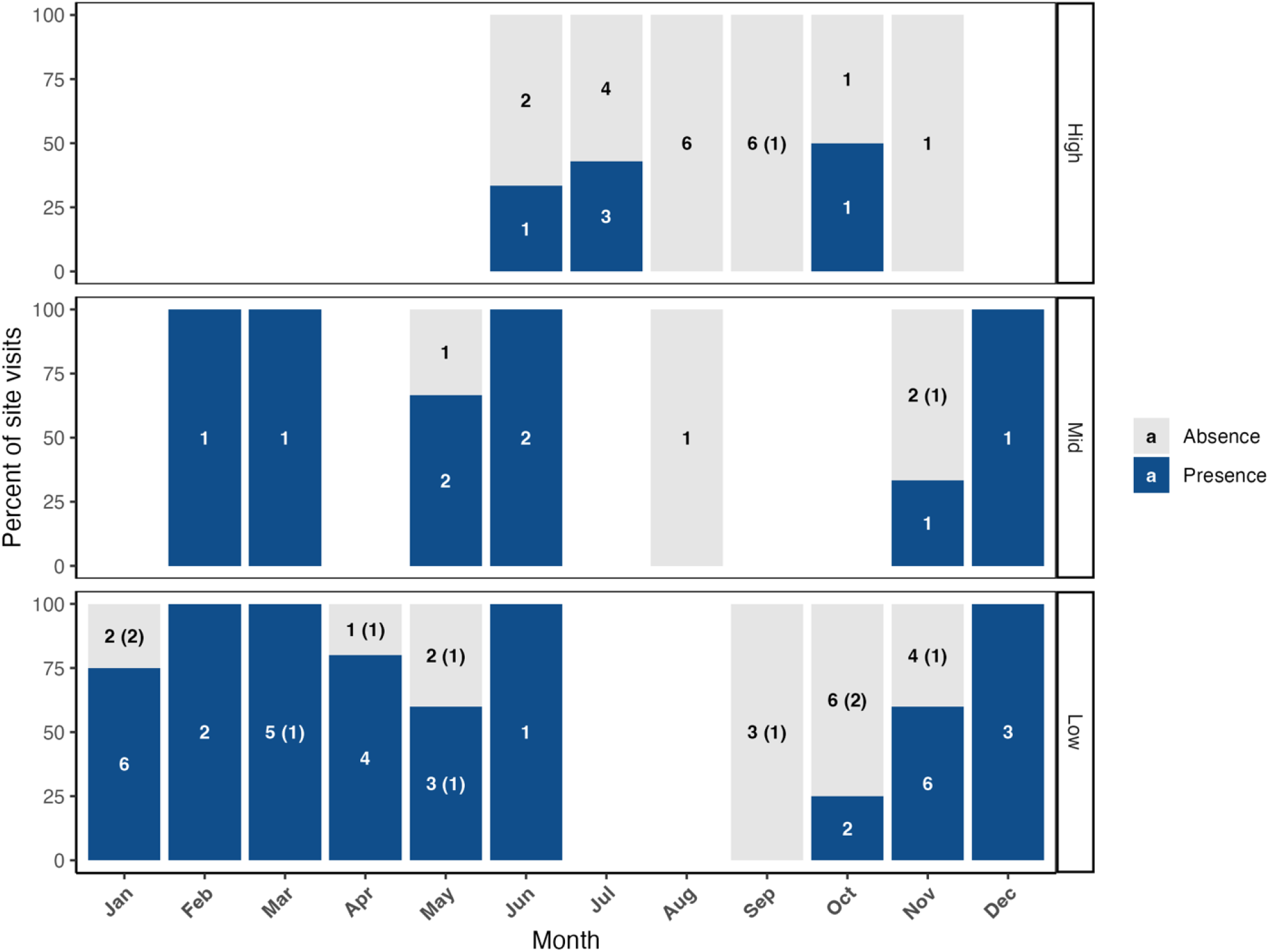
*Genea* presence in the southern Willamette Valley and associated mountains throughout a single year. Numbers within the bars indicate the number of surveys where *Genea* was found (blue) or not found (gray). Parenthetical numbers indicate how many of those sites were burned.

Odor and morphology of the truffles suggested that there are likely multiple species present. We noted that some visually similar truffles collected from shared locations had a mellow cheese odor, while others had a sharp garlic-truffle odor. Microscopic examination of these particular variants showed ellipsoid spores in the former and subglobose spores in the latter. Subsequent examinations of additional collections suggest extensive hidden diversity in the dark-colored *Genea* found in this region.

## Discussion

Using a dog trained to find truffle diversity, we found *Genea* grew abundantly nearly year-round. In the literature, *Genea* is known to be common (Alvarado et al., 2016); however, there can be an impression of local scarcity due to the difficulty of finding small and soil-like sporocarps. The commonness of *Genea* suggests that this genus may be a key ectomycorrhizal partner for trees in the area, especially the dominant Douglas-fir timber trees. Although the decrease in abundance with elevation may be partially due to a sampling bias, our findings suggest that *Genea* is most prevalent at lower elevations. Douglas-fir trees have limited distribution at higher elevations (Case & Peterson, 2005), where the landscape is more likely to be dominated by mountain hemlock (*Tsuga mertensiana*) or true firs (*Abies*), enforcing the likelihood of a strong mycorrhizal association between Douglas-fir and *Genea*.

Our surveys demonstrate that human truffle surveys have a visual bias that can be overcome by complementing with scent surveys. The dog strongly indicates, requiring the accompanying human to look more closely and find the “hidden” *Genea*. However, our surveys were limited to the abilities of the truffle dog. He cannot find truffles in hot weather, so our summer surveys were focused at higher elevations where the temperatures were cooler. Similarly, we were unable to access higher elevation sites until after snowmelt and the roads had opened, but that also would have been true of the more traditional truffle-finding method of raking.

*Genea* truffles are difficult to identify from morphological features alone, and even spores can be indistinct. Most *Genea* collections from conifer forests in the Pacific Northwest have typically been identified as *G. harknessii* (species complex) or *G. gardneri*, as the only two species of dark *Genea* currently known from this habitat (Smith et al., 2006). We have observed that there are likely several additional species that are morphologically similar and difficult to distinguish without microscopic examination and sequence data. In some instances we noted the change in odor and texture of the *Genea* we found, but we did not consistently track these traits given that these characteristics do not reliably separate different species in many hypogeous fungi. This group would benefit from further taxonomic study using modern molecular methods to determine which species are present in the area in the different habitats and elevations. However, such taxonomic study is outside the scope of this Nature Note which is focused on the ecology of the Genea genus in general and on the efficacy of using a truffle dog to find this visually indistinct truffle.

Eleven of our sites were burned, including eight conifer sites that had experienced wildfire in the past 1 to 11 years. In light of the increasing frequency and severity of wildfire in western Oregon (Dye et al., 2024; Halofsky et al., 2020), the lack of *Genea* at these sites is notable and suggests that some species of *Genea* may be sensitive to wildfire, at least in the earlier years of recovery. On the other hand, we did find *Genea* during both surveys at the burned oak savanna site, suggesting that not all *Genea* species may be equally affected by fire. This genus is relatively abundant in the Willamette Valley; however, it may decline in abundance as Douglas-fir forests are affected by climate change and high severity fire events. Our observations of *Genea* as ubiquitous in modified and disturbed habitat suggests it is resilient to anthropogenic effects, but extreme fire may be an exception for some species. We must prioritize documenting fungal species distributions, particularly ones that are difficult to locate like *Genea*, to understand what can be lost through climate change.

## Data availability statement

Data are deposited on OSF.io and will be assigned a DOI upon manuscript acceptance.

## Acknowledgements

Bitty Roy provided valuable feedback on this manuscript. This research would not have been possible without Heather’s truffle dog–good boy, Rye.

The authors have no conflicts of interest to declare.

## Notes

### Competing Interest Statement

The authors have declared no competing interest.

### Summary of Updates

Text edited and expanded following reviewer feedback

