## Supplementary material for "Use of a truffle dog provides insight into the ecology and abundant occurrence of *Genea* (Pyronemataceae) in western Oregon, U.S.A": Table S1

### Supplementary Information

**Table S1: List of sites and visiting frequency**. Note that the latitude and longitude given is for a single point, while the survey often spanned a longer distance.

| Site name | Habitat | Elevation (m) | Latitude | Longitude | Number of surveys | No. of times *Genea* found |
|---|---|---|---|---|---|---|
| Alsea Falls | Secondary Douglas-fir | 317 | 44.2944417 | -123.45409 | 1 | 1 |
| Black Creek Trail | Secondary Douglas-fir | 1115 | 43.7025533 | -122.11276 | 1 | 1 |
| Blair Lake Trail | Mature mixed conifer | 1487 | 43.8398966 | -122.24104 | 1 | 1 |
| Blue Lake | Mature mixed conifer forest | 1702 | 43.5302879 | -122.2016 | 1 | 0 |
| Blue Pool Campground | Secondary Douglas-fir | 664 | 43.7102313 | -122.29959 | 1 | 0 |
| Box Canyon Cabin | Secondary Douglas-fir | 1171 | 43.9074533 | -122.08118 | 1 | 1 |
| Canyon Creek Meadows | Mixed conifer forest | 1716 | 44.4923408 | -121.83034 | 1 | 0 |
| Carl Lake | Mixed conifer forest | 1723 | 44.5845197 | -121.78624 | 1 | 0 |
| Corrigan Lake | Lodgepole Pine | 1776 | 43.5173575 | -122.19146 | 1 | 0 |
| Creat | Secondary Douglas-fir | 333 | 43.9615467 | -123.3719 | 5 | 4 |
| Deception Creek | High severity burn (5 yrs old) | 490 | 43.7570214 | -122.55291 | 1 | 0 |
| Deception Creek | Mature secondary Douglas-fir-western hemlock forest | 399 | 43.766095 | -122.53444 | 1 | 1 |
| Divide Lake | Mature mixed conifer forest | 1526 | 49.7148559 | -119.60234 | 2 | 0 |
| Eagle's Rest | Mature secondary Douglas-fir-western hemlock forest | 812 | 43.8455167 | -122.74319 | 1 | 1 |
| Erma Bells Lake | Mature secondary Douglas-fir | 1404 | 43.8558283 | -122.04686 | 1 | 1 |
| Eugene | Urban garden | 127 | 44.0438338 | -123.12046 | 1 | 0 |
| Fairview Creek | Mixed age stand (decades to centuries) of Douglas-fir | 667 | 43.5837842 | -122.71383 | 1 | 0 |

| Site name | Habitat | Elevation (m) | Latitude | Longitude | Number of surveys | No. of times *Genea* found |
|---|---|---|---|---|---|---|
| Fall Creek | Secondary Douglas-fir | 461 | 43.98552 | -122.46043 | 1 | 1 |
| Gales Fire | High severity burn (1 yr old) | 552 | 43.9855008 | -122.43332 | 1 | 0 |
| Goodman Creek | Mature secondary Douglas-fir-western hemlock forest | 364 | 43.846905 | -122.65905 | 1 | 1 |
| Greenleaf | Secondary Douglas-fir | 169 | 44.1298 | -123.621 | 5 | 5 |
| HJ Andrews | High severity burn (2 yrs old) | 454 | 44.2040855 | -122.26005 | 1 | 0 |
| HJ Andrews | Moderate severity burn (2 yr old) | 454 | 44.2040855 | -122.26005 | 3 | 0 |
| HJ Andrews | Old growth Douglas-fir | 557 | 44.2206046 | -122.24312 | 4 | 0 |
| Hand Lake | Mature mixed conifer | 1474 | 44.2308149 | -121.87622 | 1 | 0 |
| Harlan | Secondary Douglas-fir | 113 | 44.5382083 | -123.72235 | 1 | 1 |
| [Private property] | Mature Douglas-fir | 457 | 44.183054 | -122.13557 | 1 | 0 |
| [Private property] | Secondary Douglas-fir | 457 | 44.182989 | -122.13568 | 2 | 2 |
| [Private property] east | Mature Douglas-fir | 457 | 44.1832283 | -122.13514 | 1 | 1 |
| Horse Creek Rd | Secondary Douglas-fir | 676 | 44.150275 | -122.07437 | 1 | 1 |
| Horsepasture Mtn | Mixed conifer forest | 1529 | 44.1127445 | -122.09507 | 1 | 1 |
| King Road | High severity burn (2 yrs old) | 383 | 44.155997 | -122.25237 | 1 | 0 |
| Lookout Creek | Old growth Douglas-fir | 884 | 44.23527 | -122.15383 | 1 | 0 |
| Lookout Creek | Secondary Douglas-fir | 884 | 44.23527 | -122.15383 | 1 | 1 |
| Marie and Rockpile Lakes | Mature mixed conifer forest | 1879 | 44.5519637 | -121.80433 | 1 | 0 |
| Matthieu Lakes | Mixed conifer forest | 1778 | 44.2345611 | -121.77616 | 1 | 0 |
| Matthieu Lakes | Severe burn (5 yrs) | 1778 | 44.2345611 | -121.77616 | 1 | 0 |
| Mt June | Secondary Douglas-fir | 819 | 43.816605 | -122.72048 | 1 | 1 |

| Site name | Habitat | Elevation (m) | Latitude | Longitude | Number of surveys | No. of times *Genea* found |
|---|---|---|---|---|---|---|
| Mt Pisgah | Burned oak savanna (3 yrs) | 190 | 43.996055 | -122.94983 | 2 | 2 |
| Mt Pisgah | Oak Douglas-fir savanna | 190 | 43.9972206 | -122.94442 | 1 | 1 |
| Mt Pisgah | Oak savanna | 190 | 43.9961867 | -122.94965 | 2 | 2 |
| Musick Guard Station | Secondary Douglas-fir | 1440 | 43.5813755 | -122.64143 | 1 | 1 |
| North Fork Trail | Secondary Douglas-fir | 460 | 43.7645833 | -122.48989 | 1 | 1 |
| Notch Lake | Mature mixed conifer | 1669 | 43.5789861 | -122.18071 | 1 | 0 |
| Ollalie Campground | Burn and salvage logged (~4 yr) | 711 | 44.2602533 | -122.05429 | 1 | 0 |
| Ollalie Campground | Mixed conifer forest | 711 | 44.2602533 | -122.05429 | 1 | 0 |
| Ollalie Trail | Mature Douglas-fir | 811 | 44.1494665 | -122.15536 | 1 | 0 |
| Patjens Lake Trail | High severity burn pine forest (11 yr) | 1380 | 44.3529639 | -121.89046 | 1 | 0 |
| Patjens Lake Trail | Mature mixed conifer | 1350 | 44.3495192 | -121.89685 | 1 | 0 |
| Patterson Mountain | Mixed age stand (decades to centuries) of Douglas-fir, hemlock, and fir. | 1258 | 43.7693667 | -122.61989 | 1 | 1 |
| Road to Hardesty Way | Secondary Douglas-fir | 382 | 43.662975 | -122.80781 | 1 | 1 |
| Robinson Lake | Mature mixed conifer forest | 1269 | 44.2913107 | -121.94943 | 1 | 0 |
| Rooster Rock | Secondary Douglas-fir | 495 | 44.4047105 | -122.29755 | 1 | 1 |
| Salt Creek Falls | Mixed age stand (decades to centuries) of Douglas-fir | 1287 | 43.611959 | -122.1284 | 1 | 0 |
| Sisters Mirror Lake | Mature mixed conifer forest | 1843 | 44.0402985 | -121.82812 | 1 | 0 |
| Siuslaw River Road | Secondary Douglas-fir | 194 | 43.931155 | -123.64101 | 1 | 1 |
| Starker Forest | Secondary Douglas-fir | 265 | 44.5675283 | -123.54221 | 1 | 1 |
| Tenas Lake | Mature mixed conifer forest | 1626 | 44.2292502 | -121.91689 | 2 | 0 |
| Three-Fingered Jack | Mixed conifer forest | 1716 | 44.49407 | -121.82546 | 1 | 1 |

| **Site name** | **Habitat** | **Elevation (m)** | **Latitude** | **Longitude** | **Number of surveys** | **No. of times *Genea* found** |
|---|---|---|---|---|---|---|
| Tire Mountain Trail | Mature secondary Douglas-fir | 909 | 43.8398283 | -122.5595 | 2 | 2 |
| Torrence Rd | Secondary Douglas-fir | 150 | 44.06128 | -123.45633 | 1 | 1 |
| USFS 58 | Old growth Douglas-fir | 606 | 43.799225 | -122.62547 | 1 | 1 |
| USFS 58 | Secondary Douglas-fir | 400 | 43.82853 | -122.62777 | 1 | 1 |
| Vivian Lake | Mature mixed conifer forest | 1828 | 43.5804004 | -122.16504 | 2 | 0 |
| Waldo Lake | Mature mixed conifer | 1663 | 43.6948572 | -122.04527 | 2 | 0 |
| Whittaker Creek | Secondary Douglas-fir | 153 | 43.9596333 | -123.64535 | 1 | 1 |
